# Discovering event structure in continuous narrative perception and memory

**DOI:** 10.1101/081018

**Authors:** Christopher Baldassano, Janice Chen, Asieh Zadbood, Jonathan W Pillow, Uri Hasson, Kenneth A Norman

**Author notes:** Contact: Christopher Baldassano.

## Abstract

**Summary:** During realistic, continuous perception, humans automatically segment experiences into discrete events. Using a novel model of neural event dynamics, we investigate how cortical structures generate event representations during continuous narratives, and how these events are stored and retrieved from long-term memory. Our data-driven approach enables identification of event boundaries and event correspondences across datasets without human-generated stimulus annotations, and reveals that different regions segment narratives at different timescales. We also provide the first direct evidence that narrative event boundaries in high-order areas (overlapping the default mode network) trigger encoding processes in the hippocampus, and that this encoding activity predicts pattern reinstatement during recall. Finally, we demonstrate that these areas represent abstract, multimodal situation models, and show anticipatory event reinstatement as subjects listen to a familiar narrative. Our results provide strong evidence that brain activity is naturally structured into semantically meaningful events, which are stored in and retrieved from long-term memory.

## Introduction

Typically, perception and memory are studied in the context of discrete pictures or words. Real-life experience, however, consists of a continuous stream of perceptual stimuli. The brain therefore needs to structure experience into units that can be understood and remembered: “the meaningful segments of one’s life, the coherent units of one’s personal history” (Beal & Weiss, 2013). Although this question was first investigated decades ago (Newtson, Engquist, & Bois, 1977), a general “event segmentation theory” was only proposed recently (Zacks, Speer, Swallow, Braver, & Reynolds, 2007). These and other authors have argued that humans implicitly generate event boundaries whenever the world changes in a surprising way, or when consecutive stimuli have distinct temporal associations (Schapiro, Rogers, Cordova, Turk-Browne, & Botvinick, 2013).

Two critical dimensions of event representations have not yet been deeply explored. First, events can be defined at multiple timescales. When reading a story, we could chunk it into discrete units of individual words, sentences, paragraphs, or chapters, and may need to chunk information on multiple timescales in parallel. A recent theory of cortical information processing argues for a distributed topographical hierarchy of timescales, from short processing timescales (10s to 100s of milliseconds) in early sensory regions to long processing timescales (10s to 100s of seconds) in higher-order areas (broadly overlapping the default mode network) (Hasson, Chen, & Honey, 2015). In this view, “events” in low-level sensory cortex (e.g. hearing a single word) (VanRullen, 2016) are progressively integrated into the minutes-long events typically reported by human observers.

Second, how are real-life experiences encoded into long-term memory? Behavioral experiments and mathematical models have argued that long-term memory reflects event structure during encoding (Ezzyat & Davachi, 2011; Gershman, Radulescu, Norman, & Niv, 2014; Sargent et al., 2013; Zacks, Tversky, & Iyer, 2001), suggesting that the event segments generated during perception may serve as the “episodes” of episodic memory. The well-accepted idea that the hippocampus stores “snapshots” of cortical activity has been developed with discrete memoranda (Danker, Tompary, & Davachi, 2016), for which it is obvious when snapshots should be taken and what information they should contain. However, during a continuous stream of information in a real-life context, it is not at all clear at which timescale (e.g. words, sentences, situations) snapshots should be taken, and whether these snapshots should be continuously updated during events or encoded only after an event has completed.

We propose that the full life cycle of an event, from construction to long-term storage, can be described in a unified theory, illustrated in Fig. 1. Each brain region along the processing hierarchy segments information at its preferred timescale, beginning with short events in primary visual and auditory cortex and building to multimodal, abstract representations of the features of the current event (“situation models”, Zwaan & Radvansky, 1998) in long-timescale areas, including default mode regions such as the angular gyrus and posterior medial cortex. At event boundaries in long-timescale areas, the situation model is transmitted to the hippocampus, which can later reinstate the situation model in long timescale regions during recall, and facilitate recognition of similar events in the future. This theory makes the following predictions: 1) Events should be identifiable at different timescales throughout the processing timescale hierarchy, with segmentation into short events in early sensory areas and integration into longer events in high-order areas. 2) Event boundaries annotated by human observers should be most related to neural event boundaries in long timescale regions. 3) The end of an event in long timescale cortical regions should trigger the hippocampus to encode information about the just-concluded event into episodic memory. 4) Stored event memories can be reinstated in long timescale cortical regions during recall, with stronger reinstatement for more strongly-encoded events. 5) Neural patterns in long timescale regions correspond to abstract situation models, which represent the features of the situation regardless of the way that the situation is described (e.g. a movie or a verbal narrative). 6) Prior memory for a narrative should influence the processing of future events, leading to anticipatory reinstatement in long timescale regions.

**Figure 1:**
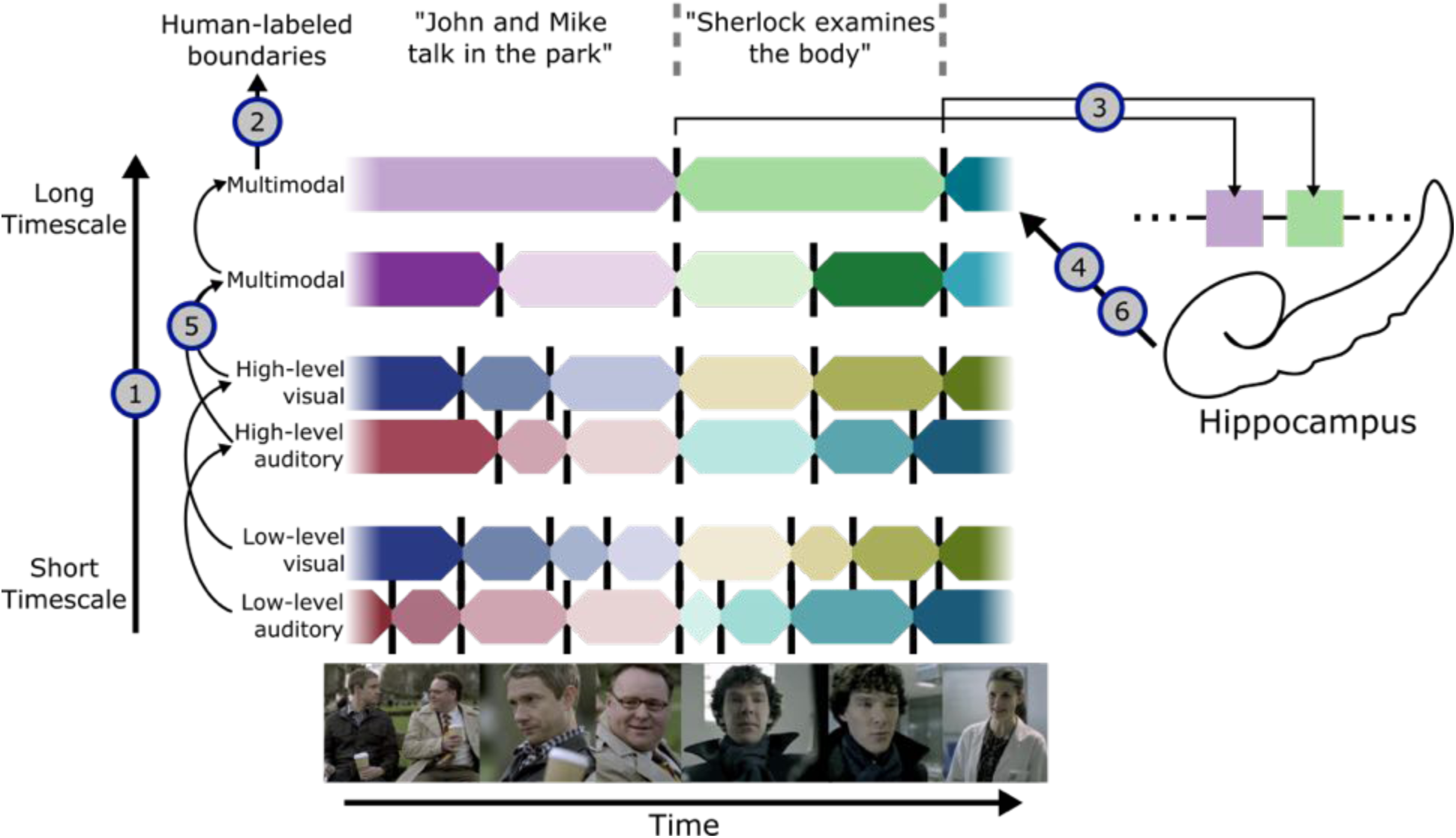
Theory of event segmentation and memory. (1) During perception, events are constructed at a hierarchy of timescales, with short events in early sensory regions (including primary visual cortex and primary auditory cortex) and long events in high-level regions (including default mode regions such as angular gyrus and posterior medial cortex). (2) Putative event boundaries identified by human observers should correspond most closely to long timescale events near the top of the hierarchy. (3) At the end of a high-level event, the situation model is stored into long-term memory, resulting in post-boundary encoding activity in the hippocampus. (4) Episodic event memories can be reinstated into high-level cortical regions during recall. (5) Since situation models are abstract semantic descriptions, the same sequence of high-level events can be activated by multiple input modalities if they describe the same story. (6) Prior event memories can also influence ongoing processing, facilitating prediction of upcoming events in related narratives. We test each of these hypotheses using a data-driven event segmentation model, which can automatically identify transitions in neural activity patterns and detect correspondences in activity patterns across datasets.

Testing this kind of integrated theory is beyond the reach of existing approaches that rely on human annotators to segment events. It requires identifying how different brain areas segment events (possibly at different timescales), and aligning events across different datasets with different timings (e.g. to see whether the same “situation model” is being elicited by a movie vs. a verbal narrative, or a movie vs. later recall). Stimulus-based annotations also cannot address questions such as anticipatory reinstatement, in which an identical stimulus generates different event segmentations in different observers (depending on their prior experience). Thus, to search for the neural correlates of event segmentation, we have developed a new data-driven analysis method that allows us to identify events directly from neural activity patterns, across multiple timescales and datasets.

Our analysis approach (summarized here, and described in detail in the *Materials and Methods*) starts with two simple assumptions: 1) while processing a particular narrative stimulus, observers progress through a particular sequence of discrete event representations (hidden states), and 2) each event has a distinct (observable) signature (a multi-voxel fMRI pattern) that is present throughout the event. We implement these assumptions using a variant of a Hidden Markov Model (HMM). Fitting the model to fMRI data (e.g. while watching a movie) entails simultaneously estimating *when* the transitions between events occur and also the mean neural pattern for each event. The optimal number of events is selected by sweeping over a range of values and maximizing the fit on held-out data. When applying the model to multiple datasets that express the same narrative (e.g. while watching a movie and during later verbal recall), the model is constrained to find the same sequence of patterns (because the events are the same), but the timing of the transitions between the patterns can vary (e.g. since the spoken description of the events might not take as long as the original events). For example, if the number of events is set to 10, the model will attempt to explain both datasets in terms of one multi-voxel pattern transitioning to a second and then a third, and so forth, where these ten patterns are common to both datasets, but exactly when the patterns switch can vary across data sets.

This model allows us to test the six predictions of our unified theory described above, following events from their initial perception in sensory cortex to their incorporation into long-term memory. Our results provide the first direct evidence that brain activity during realistic experiences is naturally structured into segmented events across multiple timescales, that event representations in high-order areas at the top of the processing hierarchy contain high-level semantic situation descriptions, and that these high-level events are discretely encoded by long-term memory structures.

## Results

All of our analyses are carried out using our new HMM-based event segmentation model (summarized above, and described in detail in the *Event Segmentation Model* subsection of *Materials and Methods*), which can automatically discover the neural signatures of each event and its temporal boundaries in a particular dataset. We validated this model using both synthetic data (Supp. Fig. 1) and narrative data with clear event breaks between stories (Supp. Fig. 2), confirming that we could accurately recover the number of event boundaries and their locations (see *Materials and Methods*). We then applied the model to test six predictions of our theory of event perception and memory.

### Timescales of cortical event segmentation

The first prediction of our theory is that “events should be identifiable at different timescales throughout the processing timescale hierarchy, with segmentation into short events in early sensory areas and integration into longer events in high-order areas.” We measured the extent to which continuous stimuli evoked the event structure hypothesized by our model (periods with stable event patterns punctuated by shifts between events), and whether the timescales of these events varied along the cortical hierarchy. We tested the model by fitting it to fMRI data collected while subjects watched a 50-minute movie (Chen, Leong, Norman, & Hasson, 2016), and then assessing how well the learned event structure explained the activity patterns of a held-out subject (by comparing within-event vs. across-event pattern similarity, with larger within- vs. across-event similarity indicating better model fit). Note that previous analyses of this dataset have shown that the evoked activity is similar across subjects, justifying an across-subjects design (Chen, Leong, et al., 2016). We found that essentially all brain regions that responded consistently to the movie (across subjects) showed evidence for event-like structure, and that the optimal number of events varied across the cortex (Fig. 2). Sensory regions like visual cortex showed faster transitions between stable activity patterns, while higher-level regions like the precuneus had activity patterns that often remained constant for over a minute before transitioning to a new stable pattern (see Fig. 2 insets). This topography of event timescales is broadly consistent with that found in previous work (Hasson et al., 2015) measuring sensitivity to temporal scrambling of a movie stimulus (see Supp. Fig. 3).

**Figure 2:**
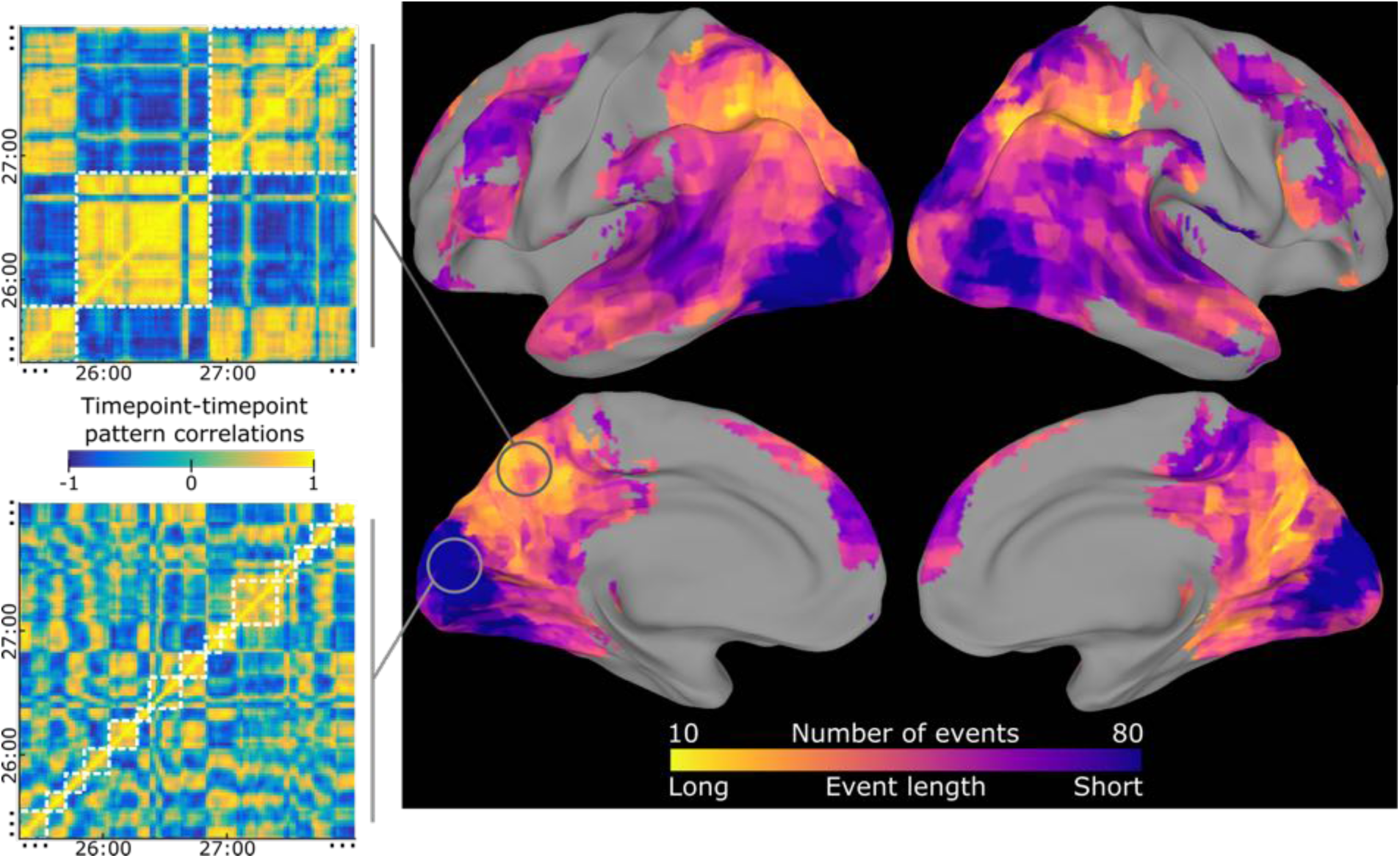
Event segmentation model for movie-watching data reveals event timescales. The event segmentation model identifies temporally-clustered structure in movie-watching data throughout all regions of cortex with high intersubject correlation. The optimal number of events varied by an order of magnitude across different regions, with a large number of short events in sensory cortex and a small number of long events in high-level cortex. For example, the timepoint correlation matrix for a region in the precuneus exhibited coarse blocks of correlated patterns, leading to model fits with a small number of events (white squares), while a region in visual cortex was best modeled with a larger number of short events (note that only ~3 minutes of the 50 minute stimulus are shown). The searchlight is masked to include only regions with intersubject correlation > 0.25, and voxelwise thresholded for greater within- than across-event similarity, q<0.001.

### Comparison to human-labeled event boundaries

Our second prediction is that “event boundaries annotated by human observers should be most related to neural event boundaries in long timescale regions.” We asked four independent raters to divide the movie into “scenes” based on major shifts in the narrative (such as in location, topic, or time). The number of event boundaries identified by the observers varied between 36 and 64, but the boundaries had a significant amount of overlap, with an average pairwise Dice’s coefficient of 0.63 and 20 event boundaries that were labeled by all four raters. We then measured, for each brain searchlight, what fraction of its neurally-defined boundaries were close to (within three timepoints of) a human-labeled event boundary. As shown in Fig. 3, this revealed a gradient from early sensory cortex to high-level long timescale regions. Early auditory and visual cortex exhibited many neural boundaries, some near boundaries marked by a human observer but also at many other times during the movie, likely due to shifts in low-level (but not high-level) features. In long timescale regions, such as angular gyrus and especially posterior medial cortex, a majority of the neurally-identified event boundaries corresponded with a boundary marked by a human observer.

**Figure 3:**
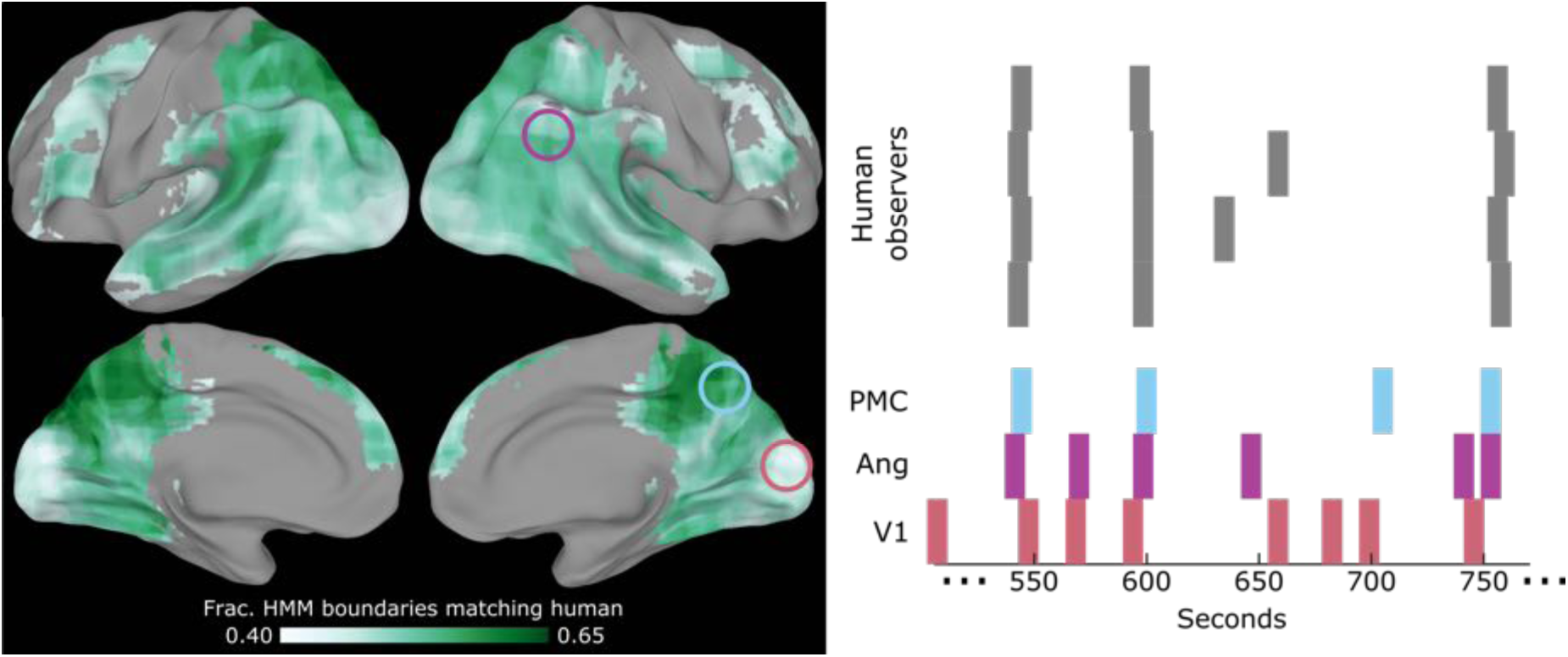
Neural event boundaries match human-labeled event boundaries, especially in posterior medial cortex. Comparing the event boundaries identified by the model to human-labeled event boundaries, we find that similarity increases as we move from sensory regions to high-level regions. The plot on the right compares human-labeled event boundaries from all four human observers to neural event boundaries for three example searchlights (for several minutes of the movie). Early sensory regions such as V1 produce a large number of boundaries that are not strongly predictive of a human-labeled event. Long timescale regions, including angular gyrus and especially in superior parietal and posterior medial cortex, have a majority of their event boundaries near human-labeled boundaries. The searchlight is masked to include only regions with intersubject correlation > 0.25.

### Relationship between cortical event boundaries and hippocampal encoding

The third prediction of our theory is that “the end of an event in long timescale cortical regions should trigger the hippocampus to encode information about the just-concluded event into episodic memory.” Prior work has shown that the end of a video clip is associated with increased hippocampal activity, and the magnitude of the activity predicts later memory (Ben-Yakov & Dudai, 2011; Ben-Yakov, Eshel, & Dudai, 2013). These experiments, however, have used only isolated short video clips with clear transitions between events. Do neurally-defined event boundaries in a continuous movie, evoked by subtler transitions between related scenes, generate the same kind of hippocampal signature? Using a searchlight procedure, we identified event boundaries with the HMM segmentation model for each cortical area across the timescale hierarchy. We then computed the average *hippocampal* activity around the event boundaries of each cortical area, to determine whether a cortical boundary tended to trigger a hippocampal response. We found that event boundaries in a distributed set of long (but not short) timescale regions (including posterior cingulate cortex and bilateral angular gyrus) all showed a strong relationship to hippocampal activity, with the hippocampal response typically peaking within several timepoints after the event boundary (Fig. 4).

**Figure 4:**
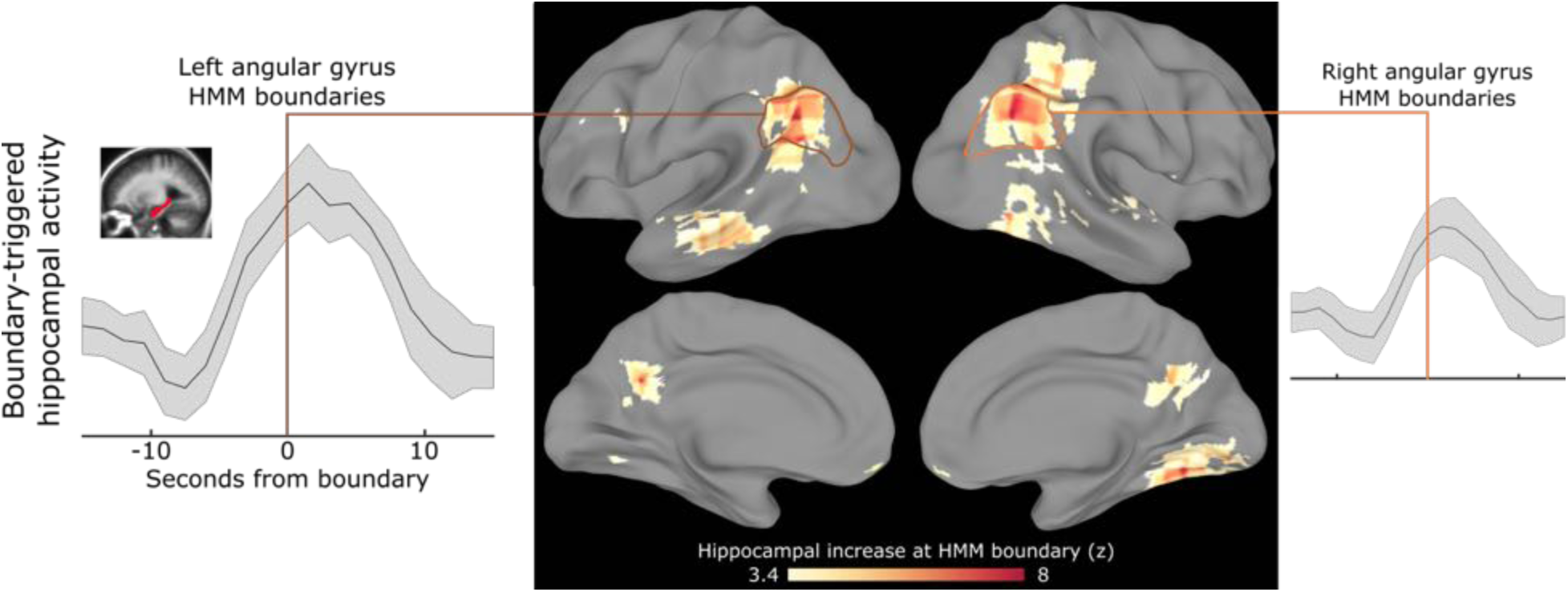
Hippocampal activity increases at cortically-defined event boundaries. To determine whether event boundaries may be related to long-term memory encoding, we identify event boundaries based on a *cortical* region and then measure *hippocampal* activity around those boundaries. In a set of high-level regions (including bilateral angular gyrus) we find that event boundaries in these regions robustly predict increases in hippocampal activity, which tends to peak just after the event boundary. The searchlight is masked to include only regions with intersubject correlation > 0.25, and voxelwise thresholded for post-boundary hippocampal activity greater than pre-boundary activity, q<0.001.

### Reinstatement of event patterns during free recall

We then tested our fourth prediction, that “stored event memories can be reinstated in long timescale cortical regions during recall, with stronger reinstatement for more strongly-encoded events.” After watching the movie, all subjects in this dataset were asked to retell the story they had just watched (without any cues or stimulus). We focused our analyses on the high-level regions that showed a strong relationship with hippocampal activity in the previous analysis (posterior cingulate and angular gyrus), as well as early auditory cortex for comparison.

Using the event segmentation model, we first estimated the (group-average) series of event-specific neural patterns evoked by the movie, and then attempted to segment each subject’s recall data into corresponding events. When fitting the model to the recall data, we assumed that the same event-specific neural patterns seen during the movie-viewing will be reinstated during the spoken recall. Analyzing the spoken recall transcriptions revealed that subjects generally recalled the events in the same order as they appeared in the movie (see table S1 in Chen, Leong, et al., 2016). Therefore, the model was constrained to use the same order of multi-voxel event patterns for recall that it had learned from the movie-watching data. However, crucially, the model was allowed to learn different event timings for the recall data compared to the movie data – this allowed us to accommodate the fact that event durations differed for free recall vs. movie-watching.

For each subject, the model attempted to find a shared sequence of latent event patterns that was shared between the movie and recall, as shown in the example with 25 events in Fig. 5a. Compared to the null hypothesis that there was no shared event order between the movie and recall, we found significant model fits in both the posterior cingulate (p=0.015) and the angular gyrus (p=0.002), but not in low-level auditory cortex (p=0.277) (Fig. 5b). This result demonstrates that we can identify shared temporal structure between perception and recall without any human annotations. A similar pattern of results can be found regardless of the number of latent events used (see Supp. Fig 4).

**Figure 5:**
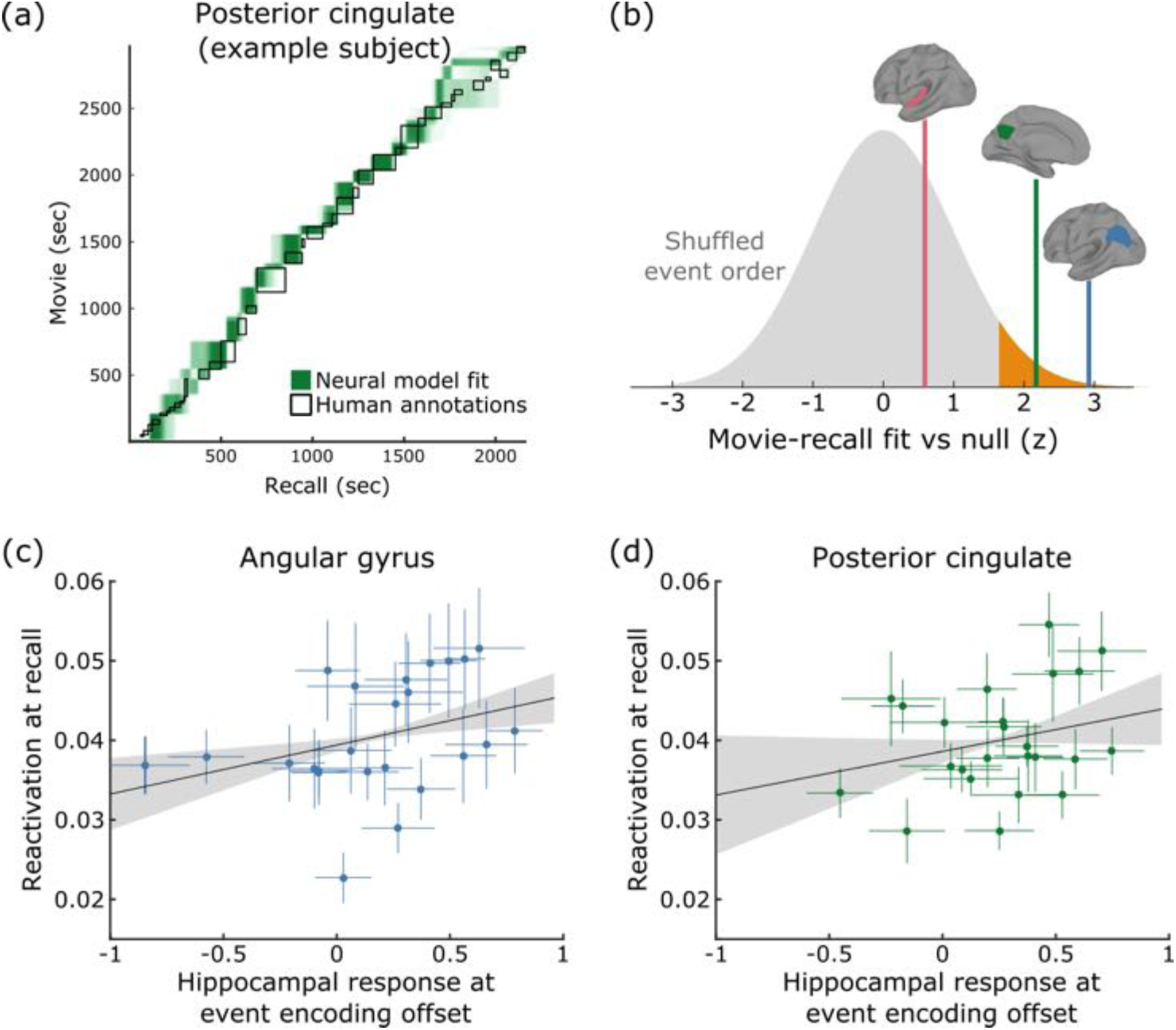
Movie-watching events are reactivated during individual free recall, and reactivation is related to hippocampal activation at encoding event boundaries. (a) We can obtain an estimated correspondence between movie-watching data and free-recall data in individual subjects by identifying a shared sequence of event patterns, shown here for an example subject using data from posterior cingulate cortex. (b) For each region of interest, we tested whether the movie and recall data shared an ordered sequence of latent events (relative to a null model in which the order of events was shuffled between movie and recall). We found that both angular gyrus (blue) and posterior cingulate cortex (green) showed significant reactivation of event patterns, while early auditory cortex (red) did not. (c-d) Events whose offset drove a strong hippocampal response during encoding (movie-watching) were strongly reactivated for longer fractions of the recall period, both in the angular gyrus and the posterior cingulate. Error bars for event points denote s.e.m. across subjects, and error bars on the best-fit line indicate 95% confidence intervals from bootstrapped best-fit lines.

We then assessed whether the hippocampal response evoked by the end of an event during the encoding of the movie to memory was predictive of the length of time for which the event was strongly reactivated during recall. As shown in Fig. 5c-d, we found that encoding activity and event reactivation were positively correlated in both angular gyrus (r=0.362, p=0.002) and the posterior cingulate (r=0.312, p=0.042), but not early auditory cortex (r=0.080, p=0.333). Note that there was no relationship between the hippocampal activity at the *starting* boundary of an event and that event’s later recall in the angular gyrus (r=−0.119, p=0.867; difference from ending boundary correlation p=0.004) and only a weak, nonsignificant relationship in posterior cingulate (r=0.189, p=0.113; difference from ending boundary correlation p=0.274).

### Shared event structure across modalities

Our fifth hypothesis is that “neural patterns in long timescale regions correspond to abstract situation models, which represent the features of the situation regardless of the way that the situation is described (e.g. a movie or a verbal narrative).” We tested this hypothesis using a separate dataset (Zadbood, Chen, Leong, Norman, & Hasson, 2016), in which some subjects watched a movie (the first 24 minutes of *Sherlock*) while other subjects listened to an 18-minute audio narration describing the events that occurred in the movie. For each cortical searchlight, we first segmented the movie data into events, and then tested whether this same sequence of events from the movie-watching subjects was present in the audio-narration subjects. High-level cortical regions with long processing timescales including the angular gyrus and posterior medial cortex showed a strongly significant correspondence between the two modalities, indicating that a similar sequence of event patterns was evoked by the movie and audio narration (Fig. 6), irrespective of the modality used to describe the events. In contrast, though low-level auditory cortex was reliably activated by both of these stimuli, there was no above-chance similarity between the series of activity patterns evoked by the two stimuli (movie vs. verbal description), presumably because the low level auditory features of the two stimuli were markedly different.

**Figure 6:**
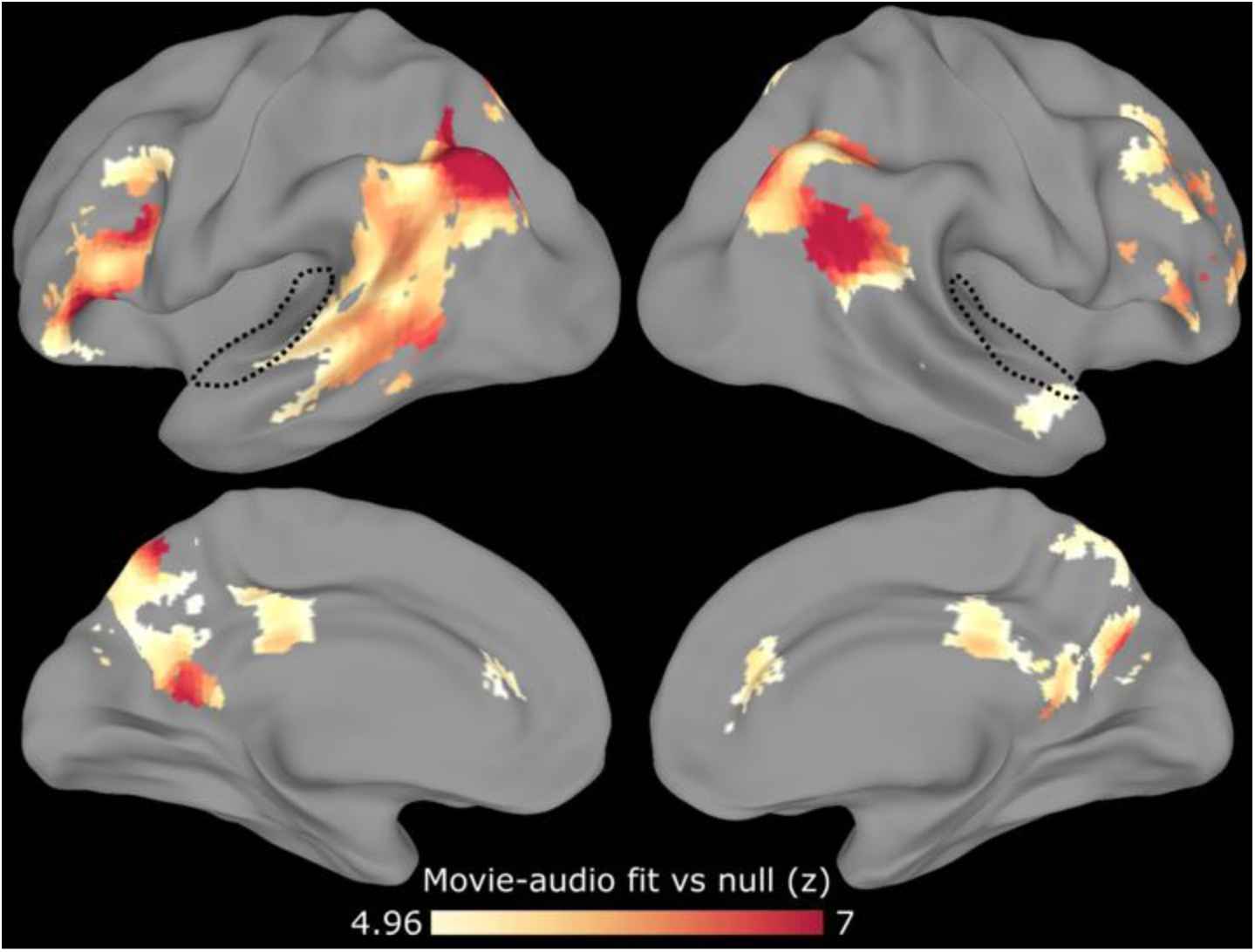
Movie-watching model generalizes to audio narration in high-level cortex. After identifying a series of event patterns in a group of subjects who watched a movie, we tested whether this same series of events occurred in a separate group of subjects who heard an audio narration of the same story. The movie and audio stimuli were not synchronized and differed in their duration. We restricted our searchlight to voxels that responded to both the movie and audio stimuli (having high ISC within each group). Movie-watching event patterns in early auditory cortex (dotted line) did not generalize to the activity evoked by audio narration, while regions including the angular gyrus and posterior medial cortex exhibited shared event structure across the two stimulus modalities. The searchlight is masked to include only regions with intersubject correlation > 0.1 in all conditions, and voxelwise thresholded for above-chance movie-audio fit, q<10^−5^.

### Anticipatory reinstatement for a familiar narrative

Finally, we tested our sixth prediction, that “prior memory for a narrative should influence the processing of future events, leading to anticipatory reinstatement in long timescale regions.” Our analyses so far have examined data on perception or on memory, but in everyday life we draw on these two functions simultaneously. Our ongoing interpretation of events can be influenced by prior knowledge; specifically, if subjects listening to the audio version of a narrative had already seen the movie version, they may anticipate upcoming events compared to subjects experiencing the narrative for the first time. Detecting this kind of anticipation has not been possible with previous approaches that rely on stimulus annotations, since the difference between the two groups is not in the stimulus (which is identical) but rather in the temporal dynamics of their cognitive processes.

We can fit our event segmentation model to the three conditions (watching the movie, listening to the narration with memory, and listening to the narration without memory) simultaneously, looking for the same sequence of event patterns in all three cases (with varying event boundaries). By analyzing which timepoints (across the three conditions) were assigned to the same event, we can generate a timepoint correspondence indicating – for each timepoint during the audio narration datasets – which timepoints of the movie are most strongly evoked (on average) in the mind of the listeners.

We searched for cortical regions along the hierarchy of timescale showing anticipation, in which this correspondence for the memory group was consistently *ahead* of the correspondence for the no-memory group (relative to chance). As shown in Fig. 7, we found anticipatory event reinstatement in several high-level regions with long processing timescales, including the angular gyrus and posterior medial cortex, with the largest leading effects in the medial frontal cortex. Examining the movie-audio correspondences in these regions, the memory group was consistently ahead of the no-memory group, indicating that for a given timepoint of the audio narration the memory group had event representations that corresponded to later timepoints in the movie.

**Figure 7:**
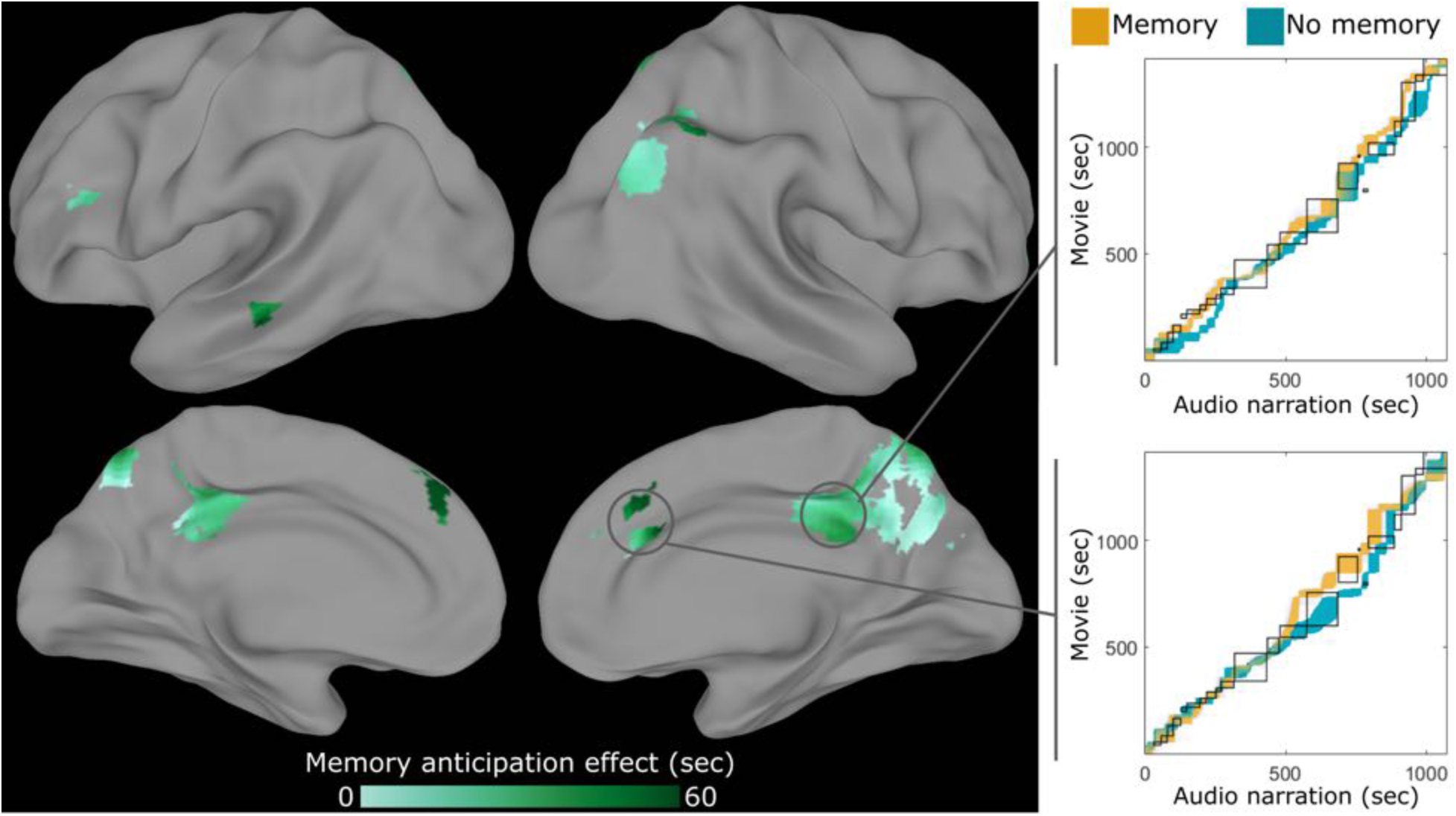
Prior memory shifts movie-audio correspondence. The event segmentation model was fit simultaneously to a data from a group watching the movie, the same group listening to the audio narration after having seen the movie (“memory”), and a separate group listening to the audio narration for the first time (“no memory”). By examining which timepoints were estimated to fall within the same latent event, we obtained a correspondence between timepoints in the audio data (for both groups) and timepoints in the movie data. We found that the correspondence in both groups was close to the human-labeled correspondence between the movie and audio stimuli (black boxes). In some regions, however, the memory correspondence (orange) significantly led the non-memory correspondence (blue), with events from the movie appearing slightly earlier for the memory group (indicated by an upward shift on the correspondence plots) despite the stimuli for the two groups being identical. This suggests that cortical regions of the memory group were anticipating events in the narration based on knowledge of the movie, with the anticipation effect increasing from posterior to anterior regions. The searchlight is masked to include only regions with intersubject correlation > 0.1 in all conditions, and voxelwise thresholded for above-chance differences between memory and no memory groups, q<0.05.

## Discussion

Using a data-driven event segmentation model that can identify temporal structure directly from neural measurements, we found that activity patterns in cortical regions including posterior medial cortex and the angular gyrus process narratives as a sequence of high-level semantic events. Although narratives evoke rapid shifts between stable activity patterns in many cortical regions along the timescale hierarchy, only these high-level regions have event representations that are closely related to human annotations, predict hippocampal encoding, are reactivated during recall, generalize across modalities, and show anticipatory coding for familiar narratives.

### Event segmentation theory

Our results are the first to demonstrate a number of key predictions of event segmentation theory (Zacks et al., 2007) directly from neural data of naturalistic narratives, without using specially-constructed stimuli or subjective labeling of where events should start and end. Previous work has shown that hand-labeled event boundaries are associated with univariate activity increases in a network of regions overlapping our high-level areas (Ezzyat & Davachi, 2011; Speer, Zacks, & Reynolds, 2007; Swallow et al., 2011; Whitney et al., 2009; Zacks, Braver, et al., 2001; Zacks, Speer, Swallow, & Maley, 2010), but by modeling fine-scale spatial activity patterns we were able to detect these event changes without an external reference. This allowed us to identify regions with temporal event structures at many different timescales, only some of which matched human-labeled boundaries. Other analyses of these datasets also found reactivation during recall (Chen, Leong, et al., 2016) and shared event structure across modalities (Zadbood et al., 2016); however, because these other analyses defined events based on the narrative rather than brain activity, they were unable to identify differences in event segmentation across brain areas or across groups with different prior knowledge.

### Timescales of perception

The topography of event timescales revealed by our analysis provides converging evidence for an emerging view of how information is processed during real-life experience (Hasson et al., 2015). The “process memory framework” argues that perceptual stimuli are integrated across longer and longer timescales along a hierarchy from early sensory regions to regions in the default mode network. Using a variety of experimental approaches, including fMRI, electrocorticography (ECoG), and single-unit recording, this topography has previously been mapped either by temporally scrambling the stimulus at different timescales to see which regions’ responses are disrupted (Hasson, Yang, Vallines, Heeger, & Rubin, 2008; Honey et al., 2012; Lerner, Honey, Silbert, & Hasson, 2011) or by examining the power spectrum of intrinsic dynamics within each region (Honey et al., 2012; Murray et al., 2014; Stephens, Honey, & Hasson, 2013). Our model and results add to these findings, by suggesting that all processing regions exhibit fast changes at event boundaries, but that these fast changes are much less frequent in long-timescale regions which accumulate and synthesize information at the situation model level, since they experience large updates only when the high-level situation model changes.

### Interactions between long timescale cortical regions and the hippocampus

Several long-timescale regions, including posterior cingulate cortex and the angular gyrus, showed effects across many of our independent analyses. These areas are involved in high-level scene processing tasks involving memory and navigation (Baldassano, Esteva, Beck, & Fei-Fei, 2016), are part of the “general recollection network” with strong anatomical and functional connectivity to the hippocampus (Rugg & Vilberg, 2013), and are the core components of the posterior medial memory system (Ranganath & Ritchey, 2012), which is thought to represent and update a representation of the current situation (Johnson-Laird, 1983; Van Dijk & Kintsch, 1983; Zwaan, Langston, & Graesser, 1995; Zwaan & Radvansky, 1998). Since event representations in these regions generalized across modalities and between perception and recall, our results provide further evidence that they encode high-level situation descriptions.

Prior work, however, has not addressed what happens to the representations in these regions when the situation changes. Behavioral experiments have shown that long-term memory reflects event structure during encoding (Ezzyat & Davachi, 2011; Sargent et al., 2013; Zacks, Tversky, et al., 2001), suggesting that situation representations are “saved” into memory as discrete events. We have demonstrated that the hippocampal encoding activity previously shown to be present at the end of movie clips (Ben-Yakov & Dudai, 2011; Ben-Yakov et al., 2013) and at abrupt switches between stimulus category and task (DuBrow & Davachi, 2016) also occurs at the much more subtle transitions between events (defined by pattern shifts in high-level regions), providing evidence that event boundaries trigger the storage of the current situation representation into long-term memory. We have also shown that this post-event hippocampal activity is related to pattern reinstatement during recall, as has been recently demonstrated for the encoding of discrete items (Danker et al., 2016), thereby supporting the view that events are the natural units of episodic memory during everyday life.

### Our event segmentation model

Temporal latent variable models have been largely absent from the field of human neuroscience, since the vast majority of experiments have a temporal structure that is defined ahead of time by the experimenter. One notable exception is the recent work of Anderson and colleagues, which has used HMM-based models to discover temporal structure in neural responses during mathematical problem solving (Anderson & Fincham, 2014; Anderson, Lee, & Fincham, 2014; Anderson, Pyke, & Fincham, 2016). These models are used to segment problem-solving operations (performed in less than 30 seconds) into a small number of cognitively distinct stages such as encoding, planning, solving and responding. Our work is the first to show that (using a modified HMM and an annealed fitting procedure) this latent-state approach can be extended to much longer experimental paradigms with a much larger number of latent states.

For finding correspondences between continuous datasets, as in our analyses of shared structure between perception and recall or perception under different modalities, several other types of approaches (not based on HMMs) have been proposed in psychology and machine learning. Dynamic time warping (Kang & Wheatley, 2015; Silbert, Honey, Simony, Poeppel, & Hasson, 2014) locally stretches or compresses two timeseries to find the best match, and more complex methods such as conditional random fields (Zhu et al., 2015) allow for parts of the match to be out of order. However, these methods do not explicitly model event boundaries, and future work will be required to investigate what types of neural correspondences are well modeled by continuous warping versus event-structured models.

### Perception and memory in the wild

Our results provide a bridge between the large literature on long-term encoding of individual items (such as words or pictures) and studies of memory for real-life experience (Nielson, Smith, Sreekumar, Dennis, & Sederberg, 2015; Rissman, Chow, Reggente, & Wagner, 2016). Since our approach does not require an experimental design with rigid timing, it opens the possibility of having subjects be more actively and realistically engaged in a task, allowing for the study of events generated during virtual reality navigation (such as spatial boundaries, Horner, Bisby, Wang, Bogus, & Burgess, 2016) or while holding dialogues with a simultaneously-scanned subject (Hasson, Ghazanfar, Galantucci, Garrod, & Keysers, 2012). The model also is not fMRI-specific, and could be applied to other types of neural timeseries such as electrocorticography (ECoG), electroencephalography (EEG), or functional near-infrared spectroscopy (fNIRS), including portable systems that could allow experiments to be run outside the lab (Mckendrick et al., 2016).

### Conclusion

Using a novel event segmentation model that can be fit directly to neuroimaging data, we showed that neural responses to naturalistic stimuli are temporally organized into discrete events at varying timescales. In a network of high-level association regions, we found that these events were related to subjective event annotations by human observers, predicted hippocampal encoding, generalized across modalities and between perception and recall, and showed anticipatory coding of familiar narratives. Our results provide a new framework for understanding how continuous experience is accumulated, stored, and recalled.

## Experimental Procedures

### Event Segmentation Model

Our model is built on two hypotheses: 1) while processing narrative stimuli, observers experience a sequence of discrete events, and 2) each event has a distinct neural signature. Mathematically, a given subject (or averaged group of subjects) starts in event s_1_=1 and ends in event s_T_=K, where T is the total number of timepoints and K is the total number of events. On each timepoint the model either remains in the current state or advances to the next state, i.e. s_t+1_ ∈ {s_t_, s_t_+1} for all timepoints t. Each event has a signature mean activity pattern m_k_ across all V voxels in a region of interest, and the observed brain activity b_t_ at any timepoint t is assumed to be highly correlated with m_k_, as illustrated in Fig. 8.

**Figure 8:**
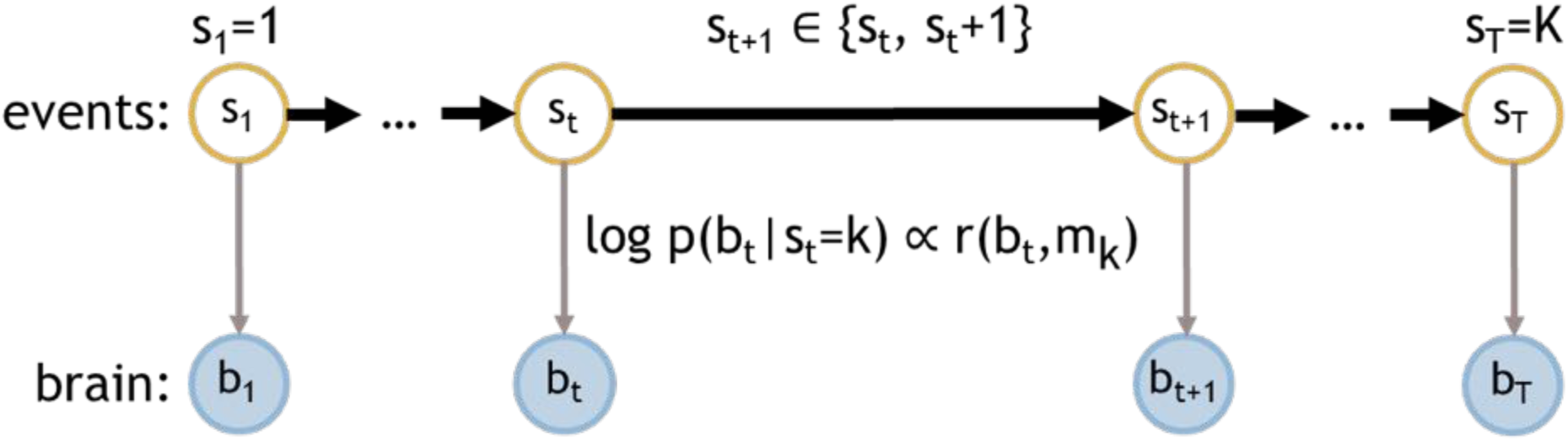
Event segmentation model. Our hypothesis about the event structure of narrative stimuli is that, for a particular story, a series of distinct events occurs in a fixed order (across stimulus modalities and between encoding and recall), and that each event k has a signature neural pattern m_k_. To encode this hypothesis in a quantitative model, we used a modified Hidden Markov Model (HMM) in which the latent state for each timepoint denotes the event to which that timepoint belongs. The model starts in the first event, and then every successive timepoint either continues the current event or starts the next event, with the final timepoint constrained to finish in the final event K. All neural datapoints during event k are assumed to be highly correlated (Pearson’s r) with m_k_.

Given the sequence of observed brain activities b_t_, our goal is to infer both the event signatures m_k_ and the event structure s_t_. To accomplish this, we cast our model as a variant of a Hidden Markov Model (HMM). The latent states are the events s_t_ that evolve according to a simple transition matrix, in which all elements are zero except for the diagonal (corresponding to s_t+1_ = s_t_) and the adjacent off-diagonal (corresponding to s_t+1_ = s_t_+1), and the observation model is an isotropic Gaussian *p*(*b*_*t*_|*s*_*t*_ = *k*) =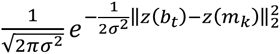, where *z*(*x*) denotes z-scoring an input vector x to have zero mean and unit variance. Note that, due to this z-scoring, the log probability of observing brain state b_t_ in an event with signature m_k_ is simply proportional to the Pearson correlation between b_t_ and m_k_ plus a constant offset.

The HMM is fit to the neural data by using an annealed version of the Baum-Welch algorithm, which iterates between estimating the neural signatures m_k_ and the latent event structure s_t_. Given the signature estimates m_k_, the event estimates p(s_t_=k) can be computed using the forward-backward algorithm. Given the event estimates p(s_t_=k), the signatures m_k_ can be computed as the weighted average 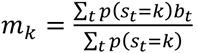. To encourage convergence to a high-likelihood solution, we anneal the observation variance *σ*^2^ as 4 ∙ 0.98^*i*^ where i is the number of loops of Baum-Welch completed so far. We stop the fitting procedure when the log-likelihood begins to decrease, indicating that the observation variance has begun to drop below the actual event activity variance. We can also fit the model simultaneously to multiple datasets; on each round of Baum-Welch, we run the forward-backward algorithm on each dataset separately, and then average across all datasets to compute a single set of shared signatures m_k_.

After fitting the model on one set of data, we can then look for the same sequence of events in another dataset. Using the signatures m_k_ learned from the first dataset, we simply perform a single round of the forward-backward algorithm to obtain event estimates p(s_t_=k) on the second dataset. If we expect the datasets to have similar noise properties (e.g. both datasets are group-averaged data from the same number of subjects), we set the observation variance to the final σ^2^ obtained while fitting the first dataset. When transferring events learned on group-averaged data to individual subjects, we estimate the variance for each event across the individual subjects of the first dataset.

The end state requirement of our model – that all states should be visited, and the end state should be symmetrical to all other states – requires extending the traditional HMM by modifying the observation probabilities *p*(*b*_*t*_|*s*_*t*_ = *k*). First, we enforce *s*_*T*_ = *K* by requiring that, on the final timestep, only the final state K could have generated the data, by setting *p*(*b*_*T*_|*s*_*T*_ = *k*) = 0 for all *k* ≠ *K*. Equivalently, we can view this as a modification of the backwards pass, by initializing the backwards message *β*(*s*_*T*_ = *k*) to 1 for (*k* = *K* and 0 otherwise. Second, we must modify the transition matrix to ensure that all valid event segmentations (which start at event 1 and end at event K, and proceed monotonically through all events) have the same prior probability. Formally, we introduce a dummy absorbing state K+1 to which state K can transition, ensuring that the transition probabilities for state K are identical to those for previous states, and then set *p*(*b*_*t*_|*s*_*t*_ = *K* + 1) to ensure that this state is never actually used. Since we do not want to assume that events will have the same relative lengths across different datasets (such as a movie and audio-narration version of the same narrative), we fix all states to have the same probability of staying in the same state (s_t+1_ = s_t_) versus jumping to the next state (s_t+1_ = s_t_+1). Note that the shared probability of jumping to the next state can take any value between 0 and 1 with no effect on the results (up to a normalization constant in the log-likelihood), since every valid event segmentation will contain exactly the same number of jumps (K-1).

Our model induces a prior over the locations of the event boundaries. There are a total of 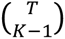 likely placements of the K-1 event boundaries, and the number of ways to have event boundary k fall on timepoint t is the number of ways that k-1 boundaries can be placed in t-1 timepoints times the number of ways that (K-1)-(k-1)-1 boundaries can be placed in T-t timepoints. Therefore *p*(*s*_*t*_ = *k* & *s*_*t*+1_ = *k* + 1)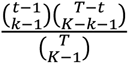. An example of this distribution is shown in Supp. Fig. 5. During the annealing process, the distribution over boundary locations starts at this prior, and slowly adjusts to match the event structure of the data.

The model implementation was first verified using simulated data. An event-structured dataset was constructed with V=10 voxels, K=10 events, and T=500 timepoints. The event structure was chosen to be either uniform (with 50 timepoints per event), or the length of each event was sampled (from first to last) from *N*(1,0.25)*(timepoints remaining)/(events remaining). A mean pattern was drawn for each event from a standard normal distribution, and the simulated data for each timepoint was the sum of the event pattern for that timepoint plus randomly distributed noise with zero mean and varying standard deviation. The noisy data were then input to the event segmentation model, and we measured the fraction of the event boundaries that were exactly recovered from the true underlying event structure. As shown in Supp. Fig. 1, were able to recover a majority of the event boundaries even when the noise level was as large as the signature patterns themselves.

Implementations of our model, along with simulated data examples, are available on github at https://github.com/intelpni/brainiak (python) and at https://github.com/cbaldassano/Event-Segmentation (Matlab).

## Experimental Data

### Interleaved Stories dataset

To test our model in a dataset with clear, unambiguous event boundaries, we used data from subjects who listened to two unrelated audio narratives (Chen, Chow, Norman, & Hasson, 2015).

22 subjects (all native English speakers) were recruited from the Princeton community (9 male, 13 female, ages 18–26). All subjects provided informed written consent prior to the start of the study in accordance with experimental procedures approved by the Princeton University Institutional Review Board. The study was approximately 2 hours long and subjects received $20 per hour as compensation for their time. Data from 3 subjects were discarded due to falling asleep during the scan, and 1 due to problems with audio delivery.

In this work we used data from 18 subjects who listened to the two audio narratives in an interleaved fashion, with the audio stimulus switching between the two narratives approximately every 60 seconds at natural paragraph breaks. The total stimulus length was approximately 29 minutes, during which there were 32 story switches. The audio was delivered via in-ear headphones.

Imaging data were acquired on a 3T full-body scanner (Siemens Skyra) with a 20-channel head coil using a T2*-weighted echo planar imaging (EPI) pulse sequence (TR 1500 ms, TE 28 ms, flip angle 64, whole-brain coverage 27 slices of 4 mm thickness, in-plane resolution 3 × 3 mm, FOV 192 × 192 mm). Preprocessing was performed in FSL, including slice time correction, motion correction, linear detrending, high-pass filtering (140 s cutoff), and coregistration and affine transformation of the functional volumes to a template brain (MNI). Functional images were resampled to 3 mm isotropic voxels for all analyses.

The analyses in this paper were carried out using data from a posterior cingulate region of interest, the posterior medial cluster in the “dorsal default mode network” defined by whole-brain resting state connectivity clustering (Shirer, Ryali, Rykhlevskaia, Menon, & Greicius, 2012).

### Sherlock Recall dataset

Our primary dataset consisted of 17 subjects who watched the first 50 minutes of the first episode of BBC’s *Sherlock*, and were then asked to freely recall the episode in the scanner without cues (Chen, Leong, et al., 2016). Subjects varied in the length and richness of their recall, with total recall times ranging from 11 minutes to 46 minutes (and a mean of 22 minutes). Imaging data was acquired using a T2*-weighted echo planar imaging (EPI) pulse sequence (TR 1500 ms, TE 28 ms, flip angle 64, whole-brain coverage 27 slices of 4 mm thickness, in-plane resolution 3 × 3 mm, FOV 192 × 192 mm).

We restricted our searchlight analyses to voxels that were reliably driven by the stimuli, measured using intersubject correlation (Hasson, Nir, Levy, Fuhrmann, & Malach, 2004). Voxels with a correlation less than r=0.25 during movie-watching were removed before running the searchlight analysis.

We defined three regions of interest based on prior work. In addition to the posterior cingulate region defined above, we defined the angular gyrus as area PG (both PGa and PGp) using the maximum probability maps from a cytoarchitectonic atlas (Eickhoff et al., 2005), and we defined early auditory cortex as voxels within the Heschl’s gyrus region (Harvard-Oxford cortical atlas) with reliable intersubject correlation during an audio narrative (“Pieman”, Simony et al., 2016).

### Sherlock Narrative dataset

To investigate cross-modal event representations and the impact of prior memory, we used a separate dataset in which subjects experienced multiple versions of a narrative. One group of 17 subjects watched the first 24 minutes of the first episode of *Sherlock* (a portion of the same episode used in the Sherlock Recall dataset), while another group of 17 subjects (who had never seen the episode before) listened to an 18 minute audio description of the events during this part of the episode (taken from the audio recording of one subject’s recall in the Sherlock Recall dataset). The subjects who watched the episode then listened to the same 18 minute audio description. This yielded three sets of data, all based on the same story: watching a movie of the events, listening to an audio narration of the events *without* prior memory, and listening to an audio narration of the events *with* prior memory. Imaging data was acquired using the same sequence as in Sherlock Recall dataset; see Zadbood et al. (2016) for full details.

As in the Sherlock Recall experiment, we removed all voxels that were not reliably driven by the stimuli. Only voxels with an intersubject correlation of at least r=0.1 across all three conditions were included in searchlight analyses.

### Event annotations by human observers

Four human observers were given the video file for the 50-minute *Sherlock* stimulus, and given the following directions: “Write down the times at which you feel like a new scene is starting; these are points in the movie when there is a major change in topic, location, time, etc. Each “scene” should be between 10 seconds and 3 minutes long. Also, give each scene a short title.” The similarity among observers was measured using Dice’s coefficient (number of matching boundaries divided by mean number of boundaries, considering boundaries within three timepoints of one another to match).

### Finding event structure in narratives

To validate our event segmentation model on real fMRI data, we first fit the model to group-averaged PCC data from the Interleaved Stories experiment. In this experiment, we expect that an event boundary should be generated every time the stimulus switches stories, giving a ground truth against which to compare the model’s segmentations. As shown in Supp. Fig. 2, our method was highly effective at identifying events, with the majority of the identified boundaries falling close to a story switch.

The remaining subsections of the *Materials and Methods* describe how the model was used to obtain each of the experimental results, with subsection titles corresponding to subsections of the *Results*.

### Timescales of cortical event segmentation

We applied the model in a searchlight to the whole-brain movie-watching data from the Sherlock Recall study. Cubical searchlights were scanned throughout the volume at a step size of 3 voxels and with a side length of 7 voxels. For each searchlight, the event segmentation model was applied to group-averaged data from all but one subject. We measured the robustness of the identified boundaries by testing whether these boundaries explained the data in the held-out subject. We measured the spatial correlation between all pairs of timepoints that were four timepoints apart, and then binned these correlations according to whether the pair of timepoints fell within the same event or crossed over an event boundary. The average difference between the within-versus across-event correlations was used as an index of how well the learned boundaries captured the temporal structure of the held-out subject. The analysis was repeated for every possible held-out subject, and with a varying number of events from K=10 to K=120. After averaging the results across subjects, the number of events with the best within-versus across-event correlations was chosen as the optimal number of events for this searchlight. To generate a null distribution, the same analysis was performed except that the event boundaries were scrambled before computing the within-versus across-event correlation. This scrambling was performed by reordering the events with their durations held constant, to ensure that the null events had the same distribution of event lengths as the real events. The within versus across difference for the real events compared to 1000 null events was used to compute a z value, which was converted to a p value using the normal distribution. The p values were Bonferroni corrected for the 12 choices of the number of events, and then the false discovery rate q was computed using the same calculation as in AFNI (Cox, 1996).

Since the topography of the results was similar to previous work on temporal receptive windows, we compared the map of the optimal number of events with the short and medium/long timescale maps derived by measuring inter-subject correlation for intact versus scrambled movies (Chen, Honey, et al., 2016). The histogram of the optimal number of events for voxels was computed within each of the timescale maps.

### Comparison to human-labeled event boundaries

To compare the neurally-defined event boundaries throughout the cortex to the human-labeled event boundaries, we computed the fraction of the neural event boundaries that were close to a human-labeled boundary for each searchlight. We defined “close to” as “within three timepoints,” since the typical uncertainty in the model about exactly where a neural event switch occurred was approximately three timepoints.

### Relationship between cortical event boundaries and hippocampal encoding

After applying the event segmentation model throughout the cortex as described above, we measured whether the data-driven event boundaries were related to activity in the hippocampus. For a given cortical searchlight, we extracted a window of mean hippocampal activity around each of the searchlight’s event boundaries. We then averaged these windows together, yielding a profile of boundary-triggered hippocampal response according to this region’s boundaries. To assess whether the hippocampus showed a significant increase in activity related to these event boundaries, we measured the mean hippocampal activity for the 10 timepoints following the event boundary minus the mean activity for the 10 timepoints preceding the event boundary, and compared this difference to the same calculation for the shuffled event boundaries (as described above). The z value for this difference was computed to a p value, and then transformed to a false discovery rate q.

### Reinstatement of event patterns during free recall

For each region of interest, we fit the event segmentation model as described above (on the group-averaged data). We then took the learned sequence of event signatures m_k_ and ran the forward-backward algorithm on each individual subject’s recall data. We set the variance of each event’s observation model by computing the variance within each event in the movie-watching data of individual subjects, pooling across both timepoints and subjects. We compared the log-likelihood of the fit to the recall data against a null model in which the event signatures were randomly re-ordered, and computed the z value of the true log-likelihood compared to 100 null shuffles, then converted to a p value. This null hypothesis test therefore assessed whether the recall exhibited *ordered* reactivation of the events identified during movie-watching. The analysis was run for 10 events to 60 events in steps of 5.

We operationalized the overall reinstatement of an event k, as ∑_*t*_ p(s_t_ = k); that is, the sum across all recall time points of the probability that the subject was recalling perceptual event k at that time point. We measured whether this per-event re-activation during recall could be predicted during movie-watching, based on the hippocampal response at the end of the event. For each subject, we computed the difference between hippocampal activity after versus before the event boundary as above. We then averaged the event re-activation and hippocampal offset response across subjects, and measured their correlation. To assess the robustness of these correlations, we performed a bootstrap test, in which we resampled subjects (with replacement, yielding 17 subjects as in the original dataset) before taking the average and computing the correlation. The p value was defined as the fraction of 1000 resamples that yielded correlations greater than zero. For comparison purposes, we also performed the same analysis but with hippocampal differences at the *beginning* of each event, rather than the end.

### Shared event structure across modalities

To determine whether audio narration of a story elicited the same sequence of events as a movie of that story, we used an approach similar to that used for detecting reactivation at recall. After fitting the event segmentation model to a searchlight of movie-watching data from the Sherlock Narration experiment, we took the learned event signatures m_k_ and used them to run the forward-backward algorithm on the audio narration data. Since both the movie and audio data were averaged at the group level, they should have similar levels of noise, and therefore we simply used the fit movie variance σ^2^ for the observation variance. As above, we compared to a null model in which the order of the event signatures was shuffled before fitting to the narration data, which yielded a z value that was converted to a p value and then corrected to a false discovery rate q.

### Anticipatory reinstatement for a familiar narrative

To determine whether memory changed the event correspondence between the movie and narration, we then fit the segmentation model simultaneously to group-averaged data from the movie-watching condition, audio narration no-memory condition, and audio narration with memory condition, yielding a sequence of events in each condition with the same neural signatures. We computed the correspondence between the movie states s_m,t_ and the audio no-memory states s_anm,t_ as as *p*(*s*_*m,t_1_*_ = *s*_*anm,t_2_*_ = ∑_k_*p*(*s*_*m,t_1_*_ = *k* ⋅ *p*(*s*_*anm,t_2_*_ = *k*), and similarly for the audio memory states s_am,t_. To determine if this correspondence was significantly different between the memory and no-memory conditions, we created null groups by averaging together a random half of the no-memory subjects with a random half of memory subjects, and then averaging together the remaining subjects from each group, yielding two group-averaged timecourses whose correspondences should differ only by chance. For both the real and null correspondences, we computed the differences between the group correspondences as 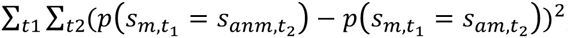, and calculated a z value based on the results for real versus null groups. This z value was converted to a p value and then corrected to a false discovery rate q. For visualization, we also computed how far the memory correspondence was ahead of the no-memory correspondence as the mean over t_2_ of the difference in the expected values 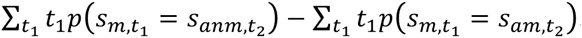.

## Acknowledgements

We thank M. Chow for assistance in collecting the Interleaved Stories dataset, M.C. Iordan for help in porting the model implementation to python, and the members of the Hasson and Norman labs for their comments and support. This work was supported by a grant from Intel Labs (CAB), The National Institutes of Health (R01-MH094480, UH, and 2T32MH065214-11, KAN), the McKnight Foundation (JWP), NSF CAREER Award IIS-1150186 (JWP), and a grant from the Simons Collaboration on the Global Brain (JWP).

## Supplementary Figures

**Supplementary Figure 1:**
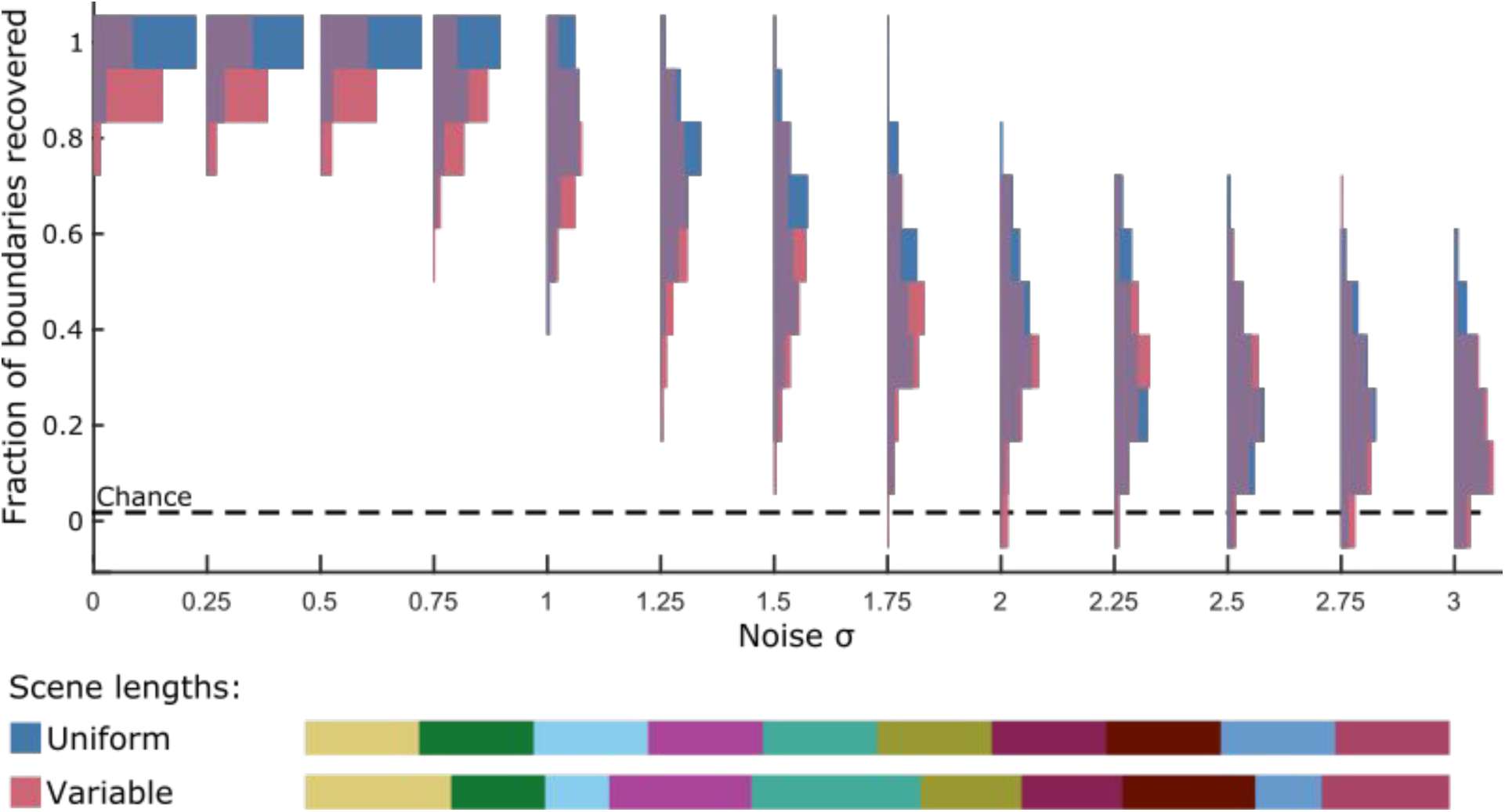
The event segmentation model recovers event boundaries on simulated data. Simulated data with a discrete event structure obscured by varying levels of noise was input to the segmentation model, with T=500, K=10, and V=10. The model successfully recovers a majority of the underlying event boundaries at low noise levels, and can still identify an above-chance fraction of boundaries even at high noise levels that are as large as the differences between the event patterns. Having variable event lengths leads to only a small loss in performance, and does not change the overall performance curve.

**Supplementary Figure 2:**
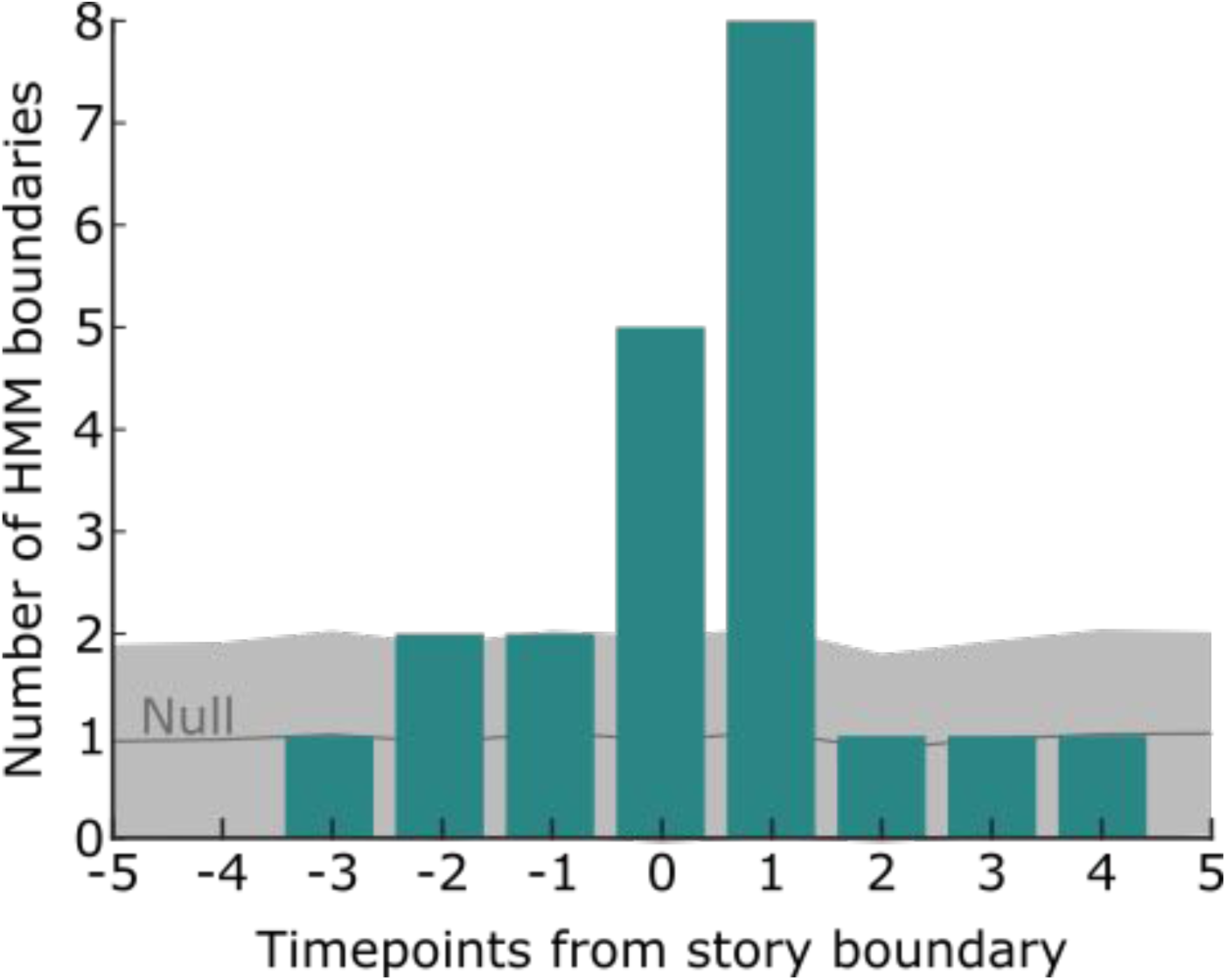
The event segmentation model successfully identifies switches between stories. Subjects listened to two stories, which were interleaved such that they alternated back and forth about every minute. Using data from PCC, an event segmentation model with 34 event transitions showed the best fit to held-out subjects (very close to the actual number of 32). Fitting the model with 34 transitions, the majority (20) were within 3 timepoints of a story switch. A null distribution was created by permuting the order of the events (preserving event lengths); under this null distribution the chance of having this many event boundaries close to true story switches was p<0.001.

**Supplementary Figure 3:**
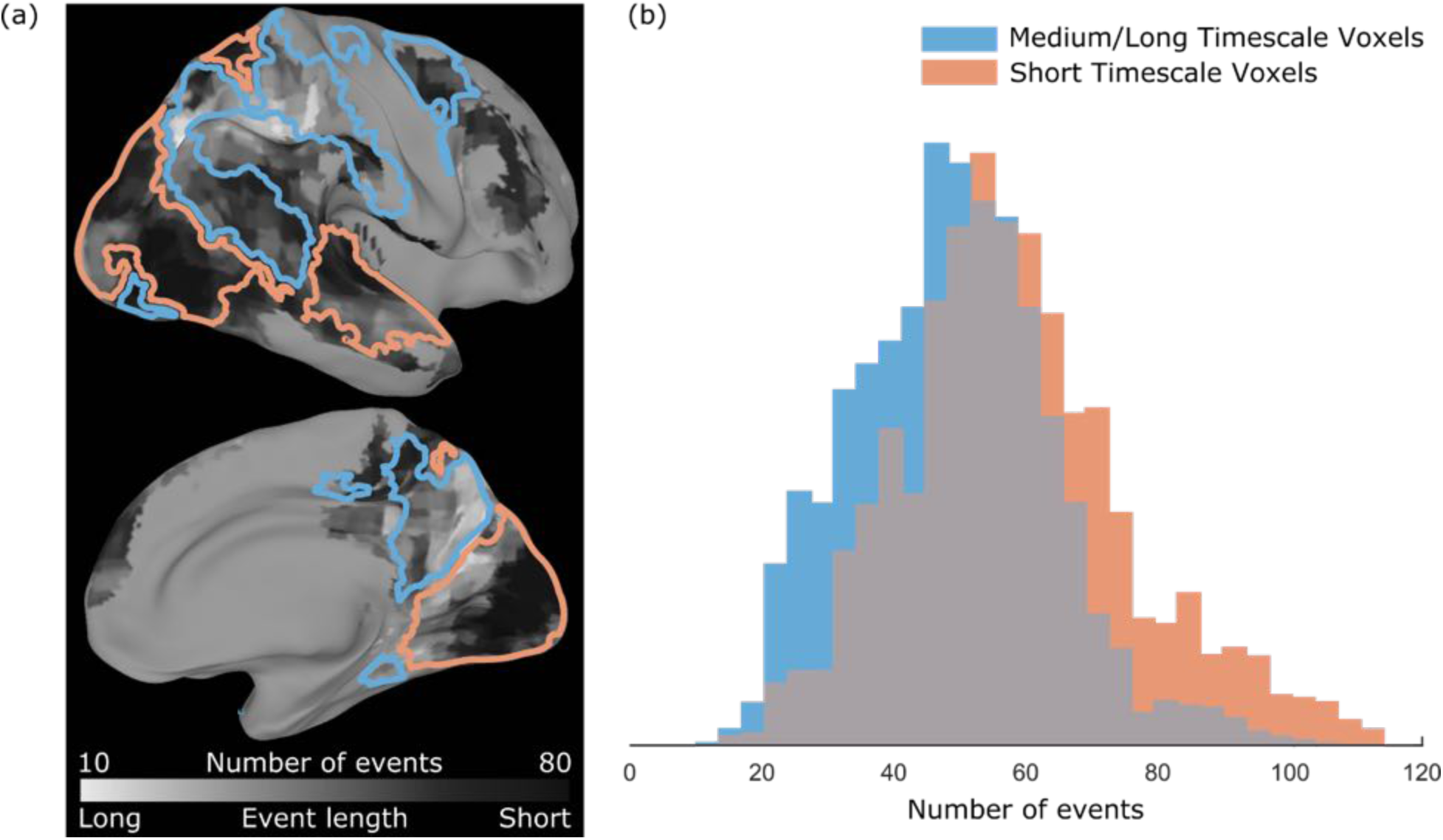
Topography of event timescales broadly matches the topography of temporal receptive windows. (a) The optimal number of events during movie watching (from Fig. 2) was compared to the map of voxel timescales (Chen, Honey, et al., 2016), which was defined based on sensitivity to temporal scrambling of a movie. Although derived from very different types of experimental data, these two approaches yield similar topographies, with early visual and auditory regions exhibiting a large number of events and having short timescales (orange), and higher-level regions having a small number of events and medium/long timescales (blue). (b) Plotting the distributions of the number of events within the short and medium/long timescale masks confirms that most regions with a small number of events have medium/long temporal receptive windows, which most regions with a large number of events have short temporal receptive windows.

**Supplementary Figure 4:**
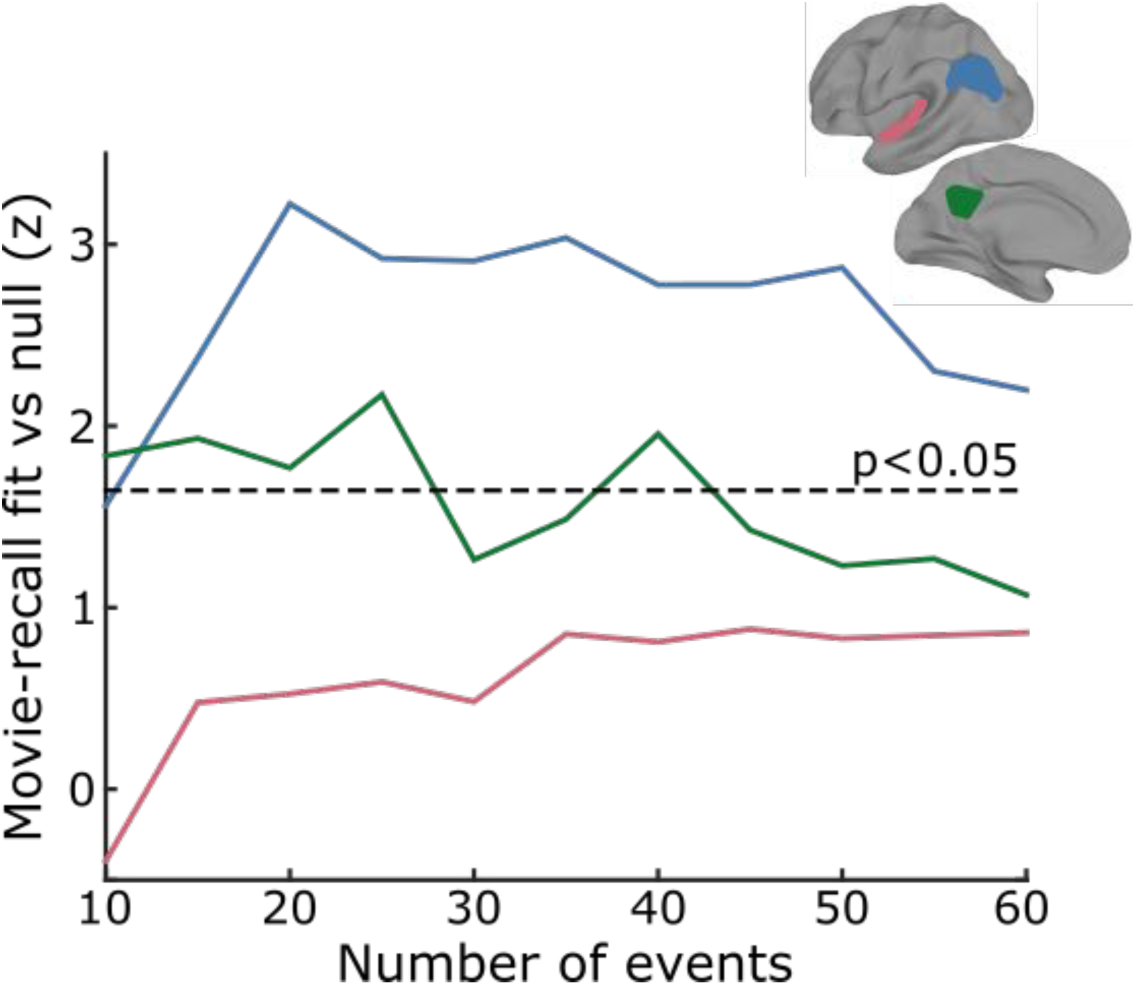
Movie and recall data show matching event structure in high-level regions across a range of settings for the number of latent events. The results shown in Fig. 5b hold for most choices of the number of latent events between 10 and 40, with decreasing goodness-of-fit for larger numbers of events. Note that the best fits were achieved with models having approximately 20-25 events, similar to the minimum number of human-labeled events that were recalled by the subjects (24, see table S1 in Chen, Leong, et al., 2016).

**Supplementary Figure 5:**
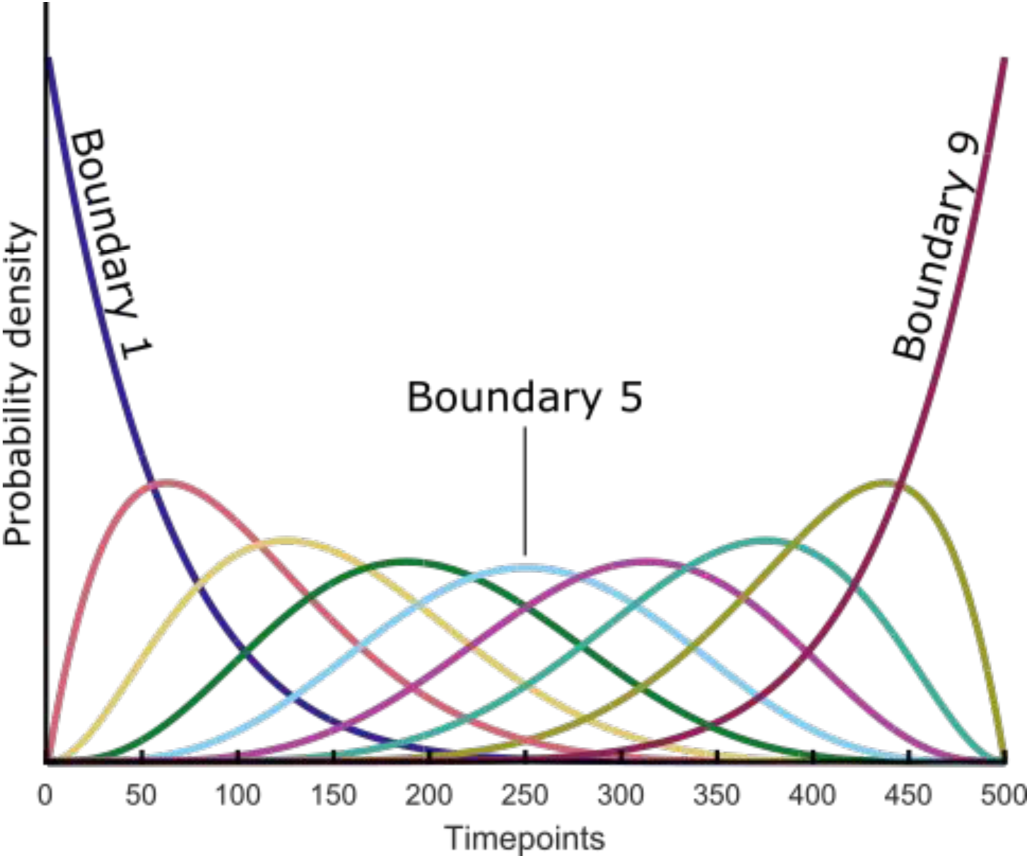
Prior distribution over event boundaries. Our event segmentation model defines a uniform prior over all possible event segmentations in which every event occurs for at least one timepoint and all events occur in order. This induces a prior distribution over event boundaries, shown here for T=500, K=10. During the annealing process, the distribution of boundaries starts at this prior, which allows for a (highly uncertain) first estimate of the signature neural pattern for each event. Based on these patterns, the latent events for all timepoints are refit, and then the patterns are recalculated. The process continues, with the pattern variance slowly decreasing, until the log likelihood reaches a peak.

